# Processing negative words in pun-humor: dynamic representation of mixed feelings blending amusement and negativity

**DOI:** 10.1101/2024.11.25.625118

**Authors:** Yingying Sun, Zhufang Jiang, Xinyue Yin, Xiaoqing Li, Ruohan Chang

## Abstract

This study aims to investigate whether amusement and negativity counteract each other or are jointly intensified during the online processing of negative keywords in pun-humor sentences, and how mixed feelings blending these two emotional states are dynamically experienced over time. Participants read three types of sentences that included the same negative words as keywords: pun-humor (negative words can generate humorous effects by resonating with the context), non-humor (negative words seamlessly align with the context), and nonsensical (negative words cannot be integrated with the context) sentences. The behavioral ratings revealed that compared with non-humor and nonsensical sentences, pun-humor sentences evoked stronger amusement and intensified negative feelings, indicating that pun-humor sentences containing negative keywords can effectively elicit mixed feelings that encompass conflicting emotional states. The ERP results showed that the N400 (300-500 ms) elicited by negative words in pun-humor sentences was comparable to that in non-humor sentences, suggesting that negative words were connected contextually in both sentence types. Besides, negative words elicited greater LPC (600-800 ms) in pun-humor sentences than in non-humor and nonsensical sentences, suggesting that pun-humor sentences require additional semantic processing and more elaborate emotional processing. Moreover, the representational similarity analysis (RSA) results revealed that in pun-humor sentences, the representation of negativity persisted for a longer duration, and it occurred and peaked earlier than that of amusement. This implies that within the dynamic representation of mixed feelings, negativity was experienced first, whereas amusement was subsequently felt within a brief period, during which negativity was not offset but rather continued to be represented over a longer time span, resulting in the simultaneous presence of both amused and negative feelings. Taken together, these findings show that mixed feelings elicited during the online processing of negative keywords in pun-humor sentences can be dynamically experienced in a way that aligns with the highly simultaneous pattern and that negativity can be experienced before amusement during the dynamic representation of mixed feelings.

---

In human social interactions, humor plays a crucial role as it fascinatingly connects unrelated elements and evokes positive emotions (Goel & Dolan, 2001; Perchtold-Stefan et al., 2020; Prenger et al., 2023). As one of the types of humor most enjoyed by recipients (Gibson & Sagarin, 2023), pun-humor effectively generates comical effects by intentionally using a keyword, which includes two possible interpretations by sharing the same phonology (homophone puns) or orthographic form (homograph puns) (Zheng & Wang, 2023a), to create ambiguity in a sentence or phrase (Koleva et al., 2019). Taking the following pun-humor, for example, *“What is critical for Jim must be hip-hop, because he is hypocritical”*. To successfully appreciate the pun-humor, one must be able to access two meanings of the keyword, its lexical salient meaning (*hypocritical*) and another plausible meaning supported by the context (*hip-hop critical*), by recognizing their shared phonology. Due to such delightful effect of certain ambiguity, pun-humor has gained widespread popularity in both literary works and advertising slogans (Zheng & Wang, 2022), and it seems that most people can find it amused (Gibson & Sagarin, 2023). However, when a pun-humor sentence incorporates a negative word as its keyword, like the example mentioned above (*hypocritical*), its online processing can not only generate amusement but also induce a sense of negativity, which may lead to a relatively complex emotional experience. During this process where negativity and amusement intertwine and fluctuate dynamically, it is important, yet under-researched, to determine whether amusement and negativity counteract each other or are jointly intensified, and how these two emotional states are experienced over time, including the patterns and relative timing of amusement and negativity.

Regarding to the first question, whether the amusement and negativity conveyed by a pun-humor sentence containing negative keyword counteract each other or are jointly intensified, existing studies and related emotion theories provide some insights. Previous studies have investigated the influence of humor on negative emotions by presenting humorous material after initial exposure to negative stimuli, and reported that amusement associated with humor can mask the negativity associated with negative stimuli (e.g., Ford & Ferguson, 2004; Fritz et al., 2017; Wu et al., 2021). In these studies, humor was used as a post hoc regulation method for offline processing of negative stimuli, and the experiences of amusement and negativity were divided according to the separate presentation of humorous and negative material. This is very different from the online processing of negative keywords in pun-humor sentences, in which both amusement and negativity can be evoked within the same stimuli. Thus, despite the observed reduction in perceived negativity during offline processing (e.g., Ford & Ferguson, 2004; Fritz et al., 2017; Wu et al., 2021), it remains uncertain whether the amusement of pun-humor reduces or intensifies the perceived negativity of negative keywords during real-time language comprehension. Answering this question can have important implications for understanding whether and to what extent we can experience two emotional states of opposing valence at the same time, a topic that is currently one of the most controversial in emotion research (Willems, 2023). Some researchers, holding a ‘bipolar’ perspective, argue that valence is an irreducible bipolar dimension of affect, and consequently precludes the simultaneous experience of positivity and negativity, the two opposite poles of the same dimension (circumplex model; Russell, 1980; Russell & Carroll, 1999; Stanisławski et al., 2021). On the other hand, some researchers, holding a ‘bivariate’ perspective, argue that the affective dimensions of positivity and negativity can be seen as two separate or independent variables, making it possible to experience two opposite emotions at the same time (evaluative space model; Cacioppo & Bernston, 1994; Larsen et al., 2001; Schimmack, 2001). And the co-experience of generally conflicting emotional states is called ‘mixed feeling’ (also referred to as “mixed valence feelings” or “mixed emotions” in some studies) —a phenomenon remained understudied until recently gaining attention (Moeller et al. 2018; Pfeifer & Pexman, 2023; Vaccaro et al., 2024). If amusement and negativity can be jointly intensified when negative keywords are processed online within pun- humor sentences, it suggests that this particular type of linguistic material can offer rich opportunities to effectively evoke and study mixed feelings. The coexistence of theoretically mutually exclusive emotional states with inconsistent valences might lead to multifaceted emotional processing of stimuli and could serve as an essential topic for understanding emotional diversity and complexity (Miyamoto et al. 2010; Rohr et al., 2016; Murray et al., 2023).

If mixed feelings blending amusement and negativity occur during the online processing of pun-humor containing negative keywords, an interesting question to further explore is what the dynamic experiential patterns of amused and negative emotional states are. This concerns whether it is best to view mixed feelings as a genuinely mixed, simultaneously positive and negative experience, or as a rapid vacillation between two different emotional states that occur so rapidly that the distinct components cannot be experienced separately (Vaccaro et al., 2020). Therefore, understanding the different patterns of mixed feelings is crucial for better addressing the abovementioned controversy between the ‘bipolar’ and ‘bivariate’ perspectives of emotion. Regarding dynamic experiential patterns, Oceja and Carrera (2009) propose the existence of at least four different patterns of mixed emotions (see Figure 1A), which can provide us with some insights and guidance for researching related issue. First, the sequential pattern: when one emotional state appears first, it is then replaced by a second emotional state, which means that the mixed feeling results from a rapid sequence of two different emotional states rather than an actual mixture of the two. Second, the inverse pattern: when the intensity of one emotional state progressively decreases, the intensity of the other progressively increases, meaning that they gradually cancel each other out during the experience until one replaces the other. Third, the prevalence pattern: when both emotional states occur simultaneously but, throughout the experiential episode, one is of moderate or high intensity while the other is very low; indeed, the latter can be considered merely a residual state and can almost be ignored, with the former being the dominant state in the mixed feeling. Finally, the highly simultaneous pattern: both emotional states are of moderate or high intensity and run a simultaneous course, either throughout the entire experiential episode or just a portion of it, indicating a period during which the two emotional states are genuinely mixed. Notably, while both the bipolar and bivariate perspectives acknowledge the existence of sequential, inverse, and prevalence patterns, only the bivariate perspective acknowledges the existence of highly simultaneous experiences (Oceja & Carrera, 2009). Thus, the key to addressing the aforementioned controversy about the possibility of a mixed feeling involving two opposing emotional states lies in distinguishing the crucial pattern around which that controversy revolves (i.e., highly simultaneous experience) from the other three patterns (i.e., sequential, prevalence, inverse). In other words, if mixed feelings can be dynamically experienced in the highly simultaneous pattern, it can provide evidence for the bivariate perspective that amusement and negativity are not mutually exclusive; when one emotional state is strong, the inhibitory effect of this emotional state is not sufficient to eliminate the theoretically opposite emotional state, meaning that people can genuinely simultaneously experience amusement and negativity.

**Figure 1.**
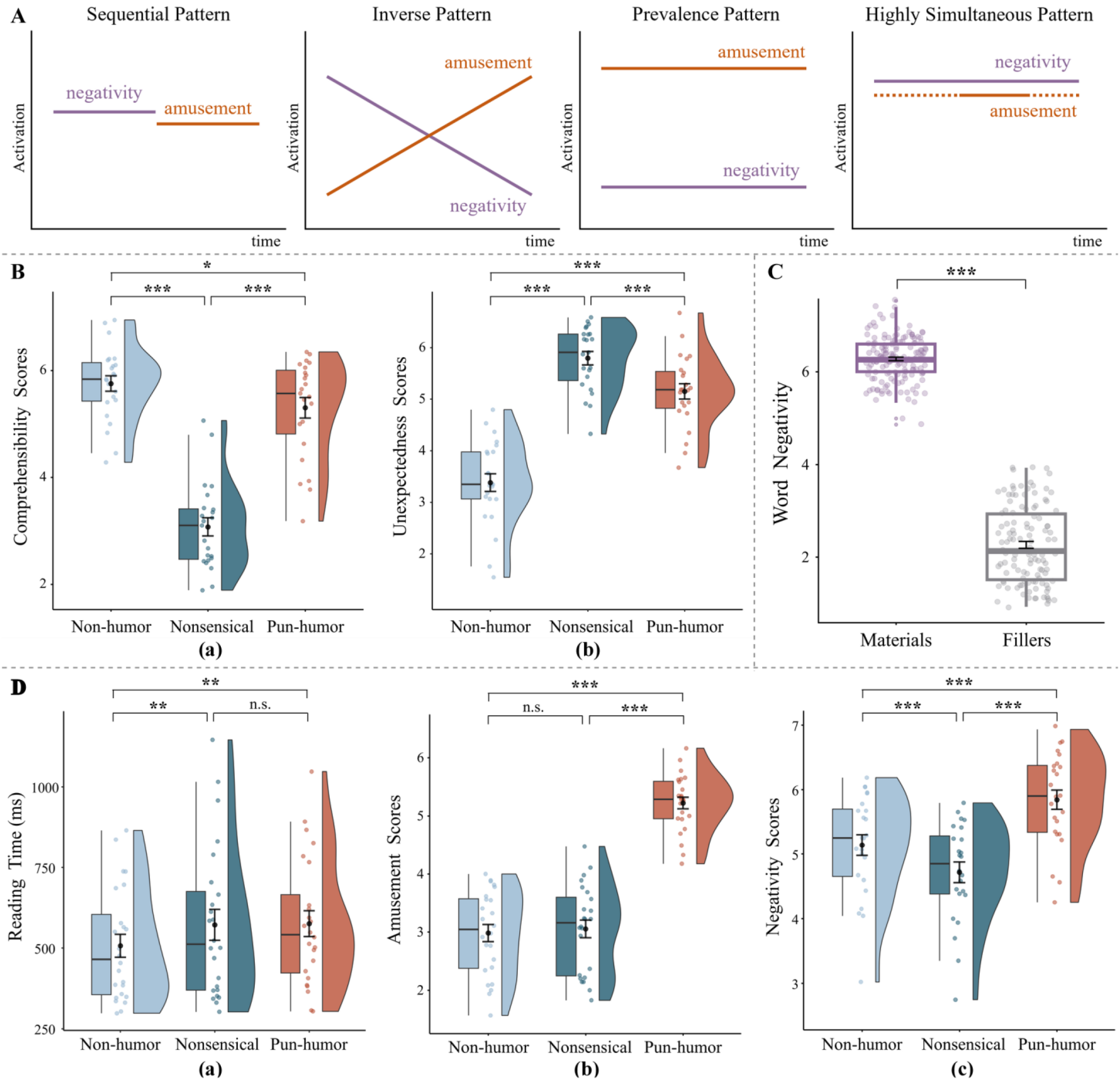
A, Diagrams of four different patterns of mixed feelings inspired by the mixed emotion patterns proposed by Oceja and Carrera (2009): Sequential Pattern, Inverse Pattern, Prevalence Pattern, Highly Simultaneous Pattern. B, The results of material evaluation before formal experiments. (a) The ratings of comprehensibility, (b) the ratings of unexpectedness for the three types of sentences. C, The ratings of negativity implied by the words used in the experimental materials and the words used in the fillers. D, The results of behavioral experiment. (a) Reading time, (b) the ratings of amusement, (c) the ratings of negativity for the three types of sentences. * = *p* < 0.05; ** = *p* < 0.01; *** = *p* < 0.001.

Moreover, beyond understanding the experiential patterns of different emotional states in mixed feelings, we also need to know the relative timing of the constituent emotional states in mixed feelings. And the temporal characteristics of amused and negative emotional states during the online processing of negative keywords in pun-humor are related to the cognitive mechanism underlying pun-humor processing. Previous pun-humor studies using non-negative keywords (e.g., Sheridan et al., 2009; Bekinschtein et al., 2011; Zheng & Wang, 2023b) suggested that successful comprehension of pun-humor involves accessing two plausible meanings of the keyword: its lexical salient meaning and another pragmatic meaning supported by the context. And the accessing and integration of these two meanings, which can evoke amusement, have been demonstrated to require additional cognitive effort. For example, a recent ERP study on Chinese homophone puns (Zheng & Wang, 2023a) reported that no significant increase in N400, yet an enhanced late positive complex (LPC) was observed in pun-humor condition relative to non-humor control. The comparable N400 in pun-humor and non-humor conditions suggests that the keywords in pun-humor condition can be contextually associated as in non-humor control (Lau et al., 2008; Kutas & Federmeier, 2011; Renoult et al., 2012). And the enhanced LPC in pun-humor was associated with increased cognitive engagement for more extensive semantic processing or reanalysis of the keyword (Coulson & Van Petten, 2002; Bakker et al., 2015; Canal et al., 2019). When the keyword in a pun-humor sentence is negative, like the pun-humor sentence mentioned earlier (e.g., “What is critical for Jim must be hip-hop, because he is hypocritical”), the keyword “hypocritical” inherently carries a negative connotation at the lexical level, and the humor is derived from the pragmatic use of this keyword within the sentence context “hip-hop critical”. This implies that accessing the lexical meaning of the keyword can trigger a sense of negativity, while finding or creating the unique connection between the keyword and the context can lead to a sense of amusement. However, although existing research results indicate that processing pun-humor involves increased cognitive effort in accessing and processing two semantic information, the relative timing of processing the lexical negative meaning and the humorous pragmatic meaning remains unclear.

Two models of language processing provide distinct hypotheses to explain this issue concerning the time course of accessing lexical and pragmatic information. One is the parallel model of language processing (Marslen-Wilson, 1987; Marslen-Wilson and Tyler, 1975; Pulvermüller et al., 2009; Strijkers et al., 2017), which assumes that different kinds of linguistic representations, including lexical semantic information and pragmatic information, can be processed synergistically at an early stage during online comprehension (Tomasello et al., 2022). According to this model, it can be speculated that the representation of negativity and amusement are both rapidly and simultaneously activated, because the comprehension of pragmatic meaning related to the onset of amusement could occur alongside the lexical semantic processes related to the onset of negativity. The other is the serial/cascade model of language processing (Friederici, 2002, 2011; Pickering and Garrod, 2004, 2013), which suggests that the onsets of different linguistic representations are processed sequentially in a cascading manner (Gao et al., 2024), with pragmatic information access emerging at a late stage of language comprehension, after other linguistic analyses such as lexical semantic processes. According to this model, it can be speculated that the onset of negativity representation prompted by lexical meaning would occur before the onset of amusement representation prompted by pragmatic comprehension. By the way, although existing EEG studies have explored the processing of pun-humor comprehension and have found that it involves additional semantic and elaborate emotional processing (Dholakia et al., 2016; Zheng & Wang, 2023a), most have focused only on traditional metrics such as ERP, making it challenging to investigate the relative timing of lexical negativity and humorous pragmatic meaning processing. This difficulty arises because such two information are intertwined during the online processing, preventing researchers from definitively determining which type of meaning or emotional state is reflected in ERP components like N400 and LPC; thus, they can primarily make indirect inferences about the cognitive significance behind these indicators. Therefore, a complementary method known as representational similarity analysis (RSA) was employed in this study. On the one hand, by calculating the correlation between behavioral patterns of different emotional states and neural patterns at various time points, RSA allows for tracking how neural patterns corresponding to the representation of specific emotional states change over time (Stokes et al., 2015; Popal et al., 2019; Xu et al., 2023). On the other hand, unlike traditional univariate analysis that compares ERP amplitudes by averaging responses across a number of items for different conditions at specific electrode sites, RSA is a multivariate analysis that simultaneously considers both the amplitude and the scalp distribution of brain waves, allowing it to access distributed information that is typically lost through averaging procedures (Żochowska et al., 2021). Combining these two methods can more intuitively and effectively reveal how specific emotional states associated with accessing different psycholinguistic information unfold over time.

Taken together, the present study had two main goals: (i) to investigate whether the perceived amusement and negativity counteract each other or are jointly intensified during the processing of pun-humor sentences with negative keywords and (ii) to examine how amusement and negativity are dynamically experienced over time during this online processing, which specifically involves the patterns and relative timing of representing amusement and negativity. To examine these questions, we will conduct behavioral and electroencephalogram (EEG) experiments and design three types of sentences that include the same negative keywords as stimuli (See Table 1 for the similar English example. Please note that this is not the actual Chinese example used in our experiment.): (1) pun-humor sentences, where the negative words generate amused effects by connecting with the context through phonetic similarity or polysemy, making the inherent negative connotations of the words intertwined with the amused effect of pun-humor; (2) non-humor sentences, where the negative words are semantically coherent and seamlessly align with the context, making them a purely negative statement; and (3) nonsensical sentences, where the negative words lack connection with the context and cannot be integrated, making their negativity mostly stem from the inherent negativity of the words. For the first goal, during the behavioral experiment, we would compare the ratings of amusement and negativity across three types of sentences, especially between pun-humor and non-humor sentences, to examine whether negativity decreases or increases in a pun-humor. For the second goal, during the EEG experiment, we would compare the ERPs to generally elucidate the online processing of negative keywords within the pun-humor sentences, and perform RSA at each time point to draw the time course of representing negativity and amusement. Thus, by using these methods, we can dissociate the dynamic representation of mixed feelings and explore the experiential pattern and relative timing of experiencing amusement and negativity.

**Table 1.** The similar English exemplar and relevant connotations of three sentence types.

| Condition | Example sentence | Pun word | Connotation |
| --- | --- | --- | --- |
| Pun-humor | What is critical for Jim must be hip-hop, because he is a **hypocritical** man. | **hypocritical** | ‘hypocritical’ has the similar sound pronunciation as ‘hip-hop critical’. |
| Non-humor | What is critical for Jim must be reputation, because he is a **hypocritical** man. |  |  |
| Nonsensical | What is critical for Jim must be dance, because he is a **hypocritical** man. |  |  |

Note: The examples used here are not from the actual experiment materials. They are similar English examples presented for demonstration purposes only. In the actual experiment, we used Chinese materials.

## Materials and Methods

### Participants

24 participants took part in the behavioral experiment (12 females, age range 18–26 years, M = 22.42 years, SD = 2.74 years). Another 39 participants, who were all right-handed, took part in the EEG experiment. 3 participants were excluded from the analysis because of excessive noise and artifacts in their EEG data. Therefore, the subsequent analyses were performed based on the remaining 36 participants (18 females, age range 19–25 years, M = 21.83 years, SD = 1.79 years).

All participants had normal or corrected-to-normal vision and were native Chinese speakers. They also reported no use of psychoactive medication, no history of neurological disorders or mental illness, and could tolerate negative words. None of them had been exposed to the experimental materials before these experiments. And they were reminded of their right to withdraw at any time during the experiment, provided written informed consent, and received compensation for their participation. This experimental protocol was approved by the Ethics Committee of Beijing Language and Culture University.

### Materials

At first, 180 two-character high-frequency Chinese words with evident negative connotations were selected from the Six Semantic Dimension Database (SSDD) (Wang et al., 2023) and the HowNet sentiment lexicon (Dong & Dong, 2003) for drafting the experimental materials, along with 120 words having positive or neutral connotations to create the fillers. Two highly skilled writers were recruited to create 180 pun-humor sentences. Each sentence included one of the selected negative words as its pun word, strategically positioned just before the final word to accommodate the EEG experimental paradigm. The pun-humor sentences were divided into two parts by a comma: the first part sets up a context by utilizing the same sound or orthographic form as the pun word, allowing participants to generate humorous effects when encountering the pun word, and the second part uses the pun word to make a negative evaluation of the protagonist in the sentence. Then, 20 participants (11 females, age range 19–26 years, M = 22.00 years, SD = 2.66 years) rated the amusement of these pun-humor sentences on a 7-point Likert scale (1 for very unamused and 7 for very amused). Based on the results, the 150 pun-humor sentences with the highest amusement scores (M = 5.21, SD = 0.61) were selected. Then, the 150 negative words in them were used as keywords to create non-humor and nonsensical sentences, both of which were of equal length and similar complexity to pun- humor sentences.

Consequently, a total of 150 sets of Chinese sentences were created as experimental materials, and each set included three types of sentences: pun-humor, non-humor, and nonsensical. These sentences were all 20 Chinese characters long, with a 10- character-long first part and a 10-character-long second part. Within each set, three sentences had completely identical second parts, namely they shared the same keyword. The only variation occurred in the first parts, leading to different effects: in pun- humor sentences, combining the first and second parts created humorous effects; in non-humor sentences, the first and second parts maintained semantic coherence; and in nonsensical sentences, understanding the combination of the first and second parts was challenging because the first parts were created by altering just one Chinese character or word from those in pun- humor sentences, which disrupted the connection between the first and second parts. The similar English examples of the three sentence types presented for demonstration purposes only are shown in Table 1.

In addition, 23 participants (12 females, age range 19–26, M = 22.39 years, SD = 2.79 years) rated on 7-point Likert scales for comprehensibility (1 for very difficult and 7 for very easy to comprehend the whole sentence) and unexpectedness (1 for not unexpected at all and 7 for very unexpected when reading the negative content in the second part after reading the first part) of all the experimental materials prior to the formal study. For the comprehensibility rating, a one-way repeated-measures ANOVA showed a significant main effect of sentence type (*F* [2, 44] = 125.16, *p* < 0.001, *η*^*2*^ = 0.85). Pairwise comparison with Bonferroni correction revealed that nonsensical sentences were significantly more difficult to comprehend (M = 3.07, SD = 0.17) than pun-humor sentences (M = 5.30, SD = 0.19, *p* < 0.001) and non-humor sentences (M = 5.76, SD = 0.14, *p* < 0.001). And pun-humor sentences were significantly less comprehensible than non-humor sentences (*p* = 0.024). For the unexpectedness rating, a one-way repeated-measures ANOVA showed a significant main effect of sentence type (*F* [2, 44] = 85.26, *p* < 0.001, *η*^*2*^ = 0.79). Pairwise comparison with Bonferroni correction showed that, after reading the first parts, the negative content in the second parts were more unexpected in nonsensical sentences (M = 5.80, SD = 0.13) compared to pun- humor sentences (M = 5.15, SD =0.15, *p* = 0.001) and non-humor sentences (M = 3.38, SD = 0.17, *p* < 0.001). And in pun- humor sentences, the negative content in the second parts were more unexpected than in non-humor sentences (*p* < 0.001). These results are shown in Figure 1B, suggesting that the experimental materials were well designed.

Since all the experimental materials used negative words as keywords, to mitigate the potential impact of sustained negativity, 120 selected words with positive or neutral connotations were used to create 40 pun-humor, 40 non-humor, and 40 nonsensical sentences as fillers. And a Latin square design was employed to create three counterbalanced lists based on 150 sets of sentences, each supplemented with the 120 fillers. Thus, a participant read 270 sentences in this study (150 experimental sentences with negative keywords and 120 fillers with positive or neutral keywords). Additionally, to ensure that the emotional valence of the keywords contained in both the experimental materials and fillers met the requirements of this study, the valence of these 270 words was rated on 9-point Likert scale (1 for very positive and 9 for very negative) by 15 participants (11 females, age range 22–27 years, M = 24.00 years, SD = 1.32 years). The results (Figure 1C) showed that the 150 words used in the experimental materials (M = 6.28, SD = 0.48) were significantly more negative than the 120 words used in the fillers (M = 2.26, SD = 0.52, *p* < 0.001).

### Paradigm

The entire experimental procedure had 2 stages: the behavioral experiment and the EEG experiment.

During the behavioral experiment, participants read and comprehended the sentences in one counterbalanced list via the self-paced reading method (SPR, reading at a natural speed to understand its meaning) and rated the amusement and negativity of each sentence. In each trial, the first part of sentence was presented in its entirety. After reading it, participants pressed the space key to move on to the second part. The second part was presented word-by-word, and participants pressed the space key to move on to the next word after reading each word, until the sentence-ending punctuation mark, a period (°), was presented. Using this method, we collected and compared reading time for the keywords. And participants rated the amusement (1 for very unamused and 7 for very amused) and negativity (1 for not negative at all and 7 for very negative) they felt after reading the entire sentence on 7-point Likert scales.

During the EEG experiment, the sentences in one counterbalanced list were pseudorandomly distributed into 6 blocks, each containing 45 sentences. The Rapid Sequential Visual Presentation (RSVP) approach was employed to better time-lock the onset of keywords. Specifically, a fixation sign “+” appeared for 500 ms to indicate the beginning of each trial. After a 500 ms blank screen, the first part of the sentence was presented until participants pressed the space key. Then, the second part of the sentence was presented word-by-word at the center of the screen, with each word appearing for 500 ms and a 300 ms blank screen interval between words. After each sentence was presented, the screen separately displayed two 9-point Likert scales, allowing participants to rate the amusement (1 for very unamused and 9 for very amused) and negativity (1 for not negative at all and 9 for very negative) of the sentence. Moreover, to avoid potential bias caused by the rating order, both within-subject and between-subject balance were implemented. That is, half of the participants rated amusement followed by negativity while the other half rated negativity followed by amusement in the first 3 blocks, and these orders were reversed in the last 3 blocks.

### EEG Recording and Preprocessing

Continuous electroencephalogram (EEG) signals were recorded using a flexible elastic cap with 64 Ag/AgCl electrodes, according to the international 10–20 system. Additionally, four electrodes (VEOU, VEOL, HEOL, and HEOR) were used to record electrooculogram (EOG) signals, which included eyeblinks, horizontal and vertical eye movements. EEG and EOG data were recorded by AC amplifier (Synamps, Neuroscan Inc.) with a band-pass filter of 0.05–100 Hz and a sampling rate of 500 Hz. The impedance levels of all electrodes were maintained below 5 kΩ throughout the experiment. During online recording, the EEG and EOG signals were referenced to the left mastoid electrode. In the offline analysis, the EEG data were re-referenced to the average of left and right mastoid electrodes.

The EEG signals were preprocessed with the EEGLAB toolbox (version 14.1.1; Delorme & Makeig, 2004) for MATLAB (The MathWorks) in the following steps. After importing into EEGLAB, all the signals were filtered with a high-pass cutoff point of 0.1 Hz and a low-pass cutoff point of 30 Hz to remove high-frequency noise and slow voltage drifts. Previous studies have shown that using filter settings in this range does not produce any distortions in RSA analysis (e.g., Wang et al., 2018; Hubbard & Federmeier, 2021; Wei et al., 2023). Then, the signals for each trial were segmented into 1000 ms periods (200 ms before and 800 ms after keyword onset), and were baseline corrected by subtracting the mean voltage of pre-stimulus signal (200-0 ms before keyword onset). It is necessary to demonstrate that the differences during the baseline interval were rigorously analyzed (SI Appendix, Figure S1) in this study, to ensure that the selection of baseline intervals does not contribute to the specific profile of subsequent ERP effects. The results showed that no significant baseline differences existed among the three sentence types (SI Appendix, Table S1), suggesting that the selected baseline interval is unlikely to introduce a bias affecting ERP response variations. Subsequently, portions of EEG epochs that contained large muscle artifacts or extreme voltage offsets were removed. And the data underwent Independent Component Analysis (ICA) to identify and remove components that were associated with eye blink and movement. The criterion for removing an ICA component encompassed evaluating the consistency between its shape, timing, and spatial location compared with the HEOG and VEOG signals. Additionally, artefacts with amplitudes exceeding ± 75 μV were removed from further analyses. On average, the remaining epochs were 139 trials per participant, and the number of remaining epochs among the pun-humor (M = 46.33, SD = 3.14), non-humor (M = 46.31, SD = 3.12), and nonsensical (M = 46.63, SD = 3.11) sentences showed no significant differences (*ps* = 1.000).

### ERP Analysis

To investigate the association between negative keywords and contexts, as well as the integration processing of negative keywords across three sentence types, the N400 and LPC components were compared in the ERP analysis. And the most classical time windows for N400 and LPC were selected, which were 300-500 ms and 600-800 ms, respectively (Han et al., 2020; Urbach et al., 2020). For each time window, single-trial analysis was performed on the EEG data via linear mixed effect models (LMEs) with the lme4 package (Bates et al., 2015) in R. Nine classical regions of interest (ROIs) were set up to examine the scalp distribution, each containing five or six electrodes (Figure 2A): left frontal (F3, F5, F7, FC3, FC5, and FT7), left central (C3, C5, CP3, CP5, and TP7), left parietal (P3, P5, P7, PO5, PO7, and O1), medial frontal (F1, FZ, F2, FC1, FCZ, and FC2), medial central (C1, CZ, C2, CP1, CPZ, and CP2), medial parietal (P1, PZ, P2, PO3, POZ, and PO4), right frontal (F4, F6, F8, FC4, FC6, and FT8), right central (C4, C6, CP4, CP6, and TP8), and right parietal (P4, P6, P8, PO6, PO8, and O2). The mean amplitudes of each ROI during the two time windows were computed and used as the dependent variables. During the analysis, sentence type and ROI were defined as fixed effects. The models were started with random intercepts for subjects and items, as well as by-subject and by-item random slopes for the effects of sentence type and ROI. To achieve model convergence, the final models were constructed using the following formula: DV ∼ Type * ROI + (1 + Type | Subject) + (1 | Item) (Zheng & Wang, 2023a). The significance of the predictors and their interactions were computed using the Mixed function from the afex package (Singmann et al., 2023) and post hoc pairwise comparisons were performed with Bonferroni correction using the emmeans function from the emmeans package (Lenth, 2022).

**Figure 2.**
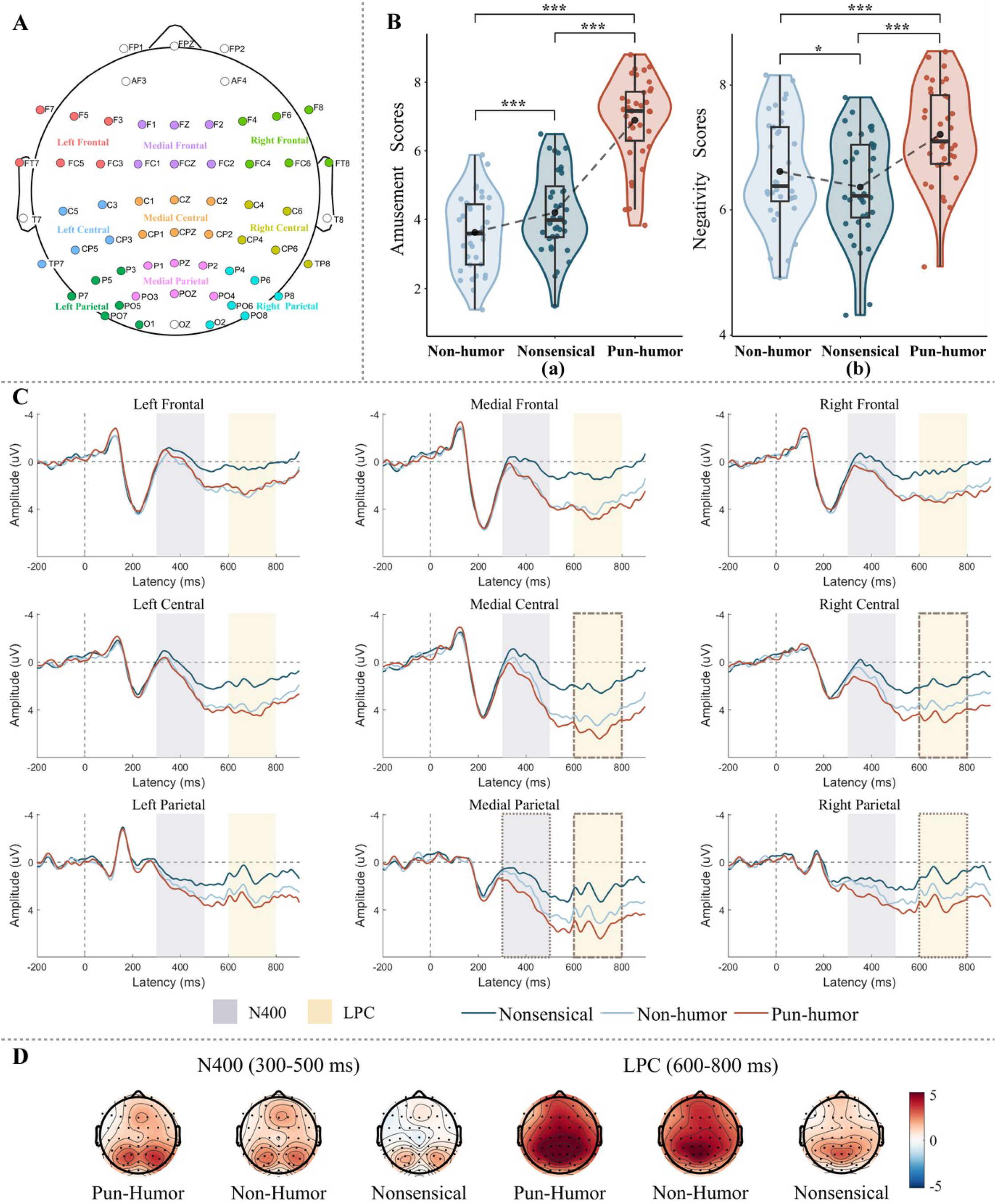
Behavioral and ERP results in EEG experiment. A, The distribution of 9 regions of interest (ROI) in our study. B, The behavioral rating results in the EEG experiment. (a) The rating results of amusement, (b) the rating results of negativity for the three types of sentences. C, The ERP results of 9 ROIs for the three types of sentences. D, The topographic maps of N400 and LPC components. * = *p* < 0.05; ** = *p* < 0.01; *** = *p* < 0.001.

### RSA analysis

To explore the relative timing of representing amusement and negativity during the online processing of negative words embedded in pun-humor sentences, we conducted RSA analysis on these sentences using the Neurora Toolbox in Python (Lu & Ku, 2020), following the analysis codes and steps of previous researchers (Gao et al., 2024). First, representational dissimilarity matrixes (RDMs) were separately constructed from behavioral data and EEG data. For behavioral RDMs, all the behavioral data were normalized to Z score, then the Euclidean distance between any two trials was calculated according to the negativity or amusement ratings of each participant. For EEG RDMs, a data vector was extracted for each trial and for each participant from a 10 ms (5 time points) sliding window with a step size of 5 time points that represented the spatial pattern of neural activity of the middle time point (t) across all 60 scalp voltage channels. At each middle time point, the dissimilarity matrices were created by calculating the pairwise dissimilarity values between the spatial patterns of neural activity of any two trials, which were quantified as (1- Pearson’s r) values between the data vectors. Next, to identify when neural representations underlying specific aspects of emotion emerge, partial correlations were calculated between the different behavioral RDMs and the EEG RDMs at each middle time point for each participant. For each emotional aspect, when calculating its partial correlation with EEG RDMs, the impact of other irrelevant variables is excluded, aiming to concentrate exclusively on the correlation between the variable under investigation and the EEG RDMs (Kato et al., 2022). Specifically, for amusement, the effects of negativity and the comprehensibility, unexpectedness, and word negativity of corresponding materials were partialled out. For negativity, the effects of amusement and the comprehensibility, unexpectedness, and word negativity of corresponding materials were partialled out. Moreover, it is important to note that only the upper triangle of each RDM was extracted as vectors to calculate these correlations (Ritchie et al., 2017), because the upper and lower triangles were mirror images of each other, and the diagonal cells always had values of 0 (Kiat et al., 2022).

Furthermore, cluster-based permutation tests were performed to evaluate the temporal clusters in which the correlations showed significant effects. The null hypothesis corresponded to a correlation coefficient of zero, while significant temporal clusters were specified as adjacent time points with statistical values exceeding the cluster-inducing threshold. To draw the t-value distribution under the null hypothesis, the permutation test was conducted by randomly shuffling the data and calculating the results for 2000 iterations, through which the statistical significance could be assessed. At each time point, the cluster-inducing threshold was set as the 95th percentile of the t-value distribution (equivalent to *p* < 0.05, one-sided), which means that real data exceeding 95% of the distribution were regarded as significant (Li et al., 2022).

Finally, the Bootstrap test was used to estimate statistical differences of the onset (i.e., the initial significant time point), peak (i.e., the time point where the peak value of correlation coefficient is observed), and duration (i.e., the length of the significant time period) latencies between the representation of negativity and amusement in pun-humor sentences. Each bootstrap sampling iteration involved drawing, with replacement, n subjects from the entire pool of subjects to form a new sample, then the latencies for this sample can be acquired. This process was repeated 1000 times, resulting in empirical distributions of the onset, peak, and duration latencies for the representation of negativity and amusement. The p-value was the number of samples whose differences between the latencies were bigger or smaller than zero divided by the number of bootstrap samples (i.e., 1000) (Li et al., 2022).

Moreover, to explore the relative timing of representing negativity during the online processing of negative words among three different sentence types, a similar RSA procedure was also conducted on non-humor and nonsensical sentences. And the Bootstrap test was conducted to estimate the statistical differences in onset and duration latencies among time course of representing negativity in the three sentence types. It should be added that these p-values were corrected for multiple comparisons with a false discovery rate (FDR) of 0.05.

## Results

### Results of the behavioral experiment

Figure 1D shows the average reading time, amusement rating, and negativity rating across the three sentence types in the behavioral experiment.

For reading time, a one-way repeated-measures ANOVA revealed a significant main effect of sentence type (*F* [2, 46] = 8.79, *p* < 0.001, *η*^*2*^ = 0.28). Pairwise comparison with Bonferroni correction revealed that participants spent significantly less time reading keywords in non-humor sentences (M = 506.98 ms, SD = 173.63 ms) than in both pun-humor (M = 575.91 ms, SD = 197.75 ms; *p* = 0.003) and nonsensical sentences (M = 572.27 ms, SD = 235.06 ms; *p* = 0.006). The difference in reading time between pun-humor and nonsensical sentences was not significant (*p* = 1.000).

For the amusement rating, a one-way repeated-measures ANOVA revealed a significant main effect of sentence type (*F* [2, 46] = 136.09, p < 0.001, *η*^*2*^ = 0.86). Pairwise comparison with Bonferroni correction revealed that pun-humor sentences (M = 5.22, SD = 0.49) were significantly more amusing than both non-humor (M = 2.98, SD = 0.72; *p* < 0.001) and nonsensical sentences (M = 3.05, SD = 0.75; *p* < 0.001), and there was no significant difference in amusement between non-humor and nonsensical sentences (*p* = 1.000).

For the negativity rating, a one-way repeated-measures ANOVA revealed a significant main effect of sentence type (*F* [2, 46] = 43.99, *p* < 0.001, *η*^*2*^ = 0.66). Pairwise comparison with Bonferroni correction revealed that pun-humor sentences (M = 5.84, SD = 0.72) were significantly more negative than both non-humor (M = 5.13, SD = 0.78; *p* < 0.001) and nonsensical sentences (M = 4.71, SD = 0.76; *p* < 0.001), and non-humor sentences were significantly more negative than nonsensical sentences (*p* < 0.001).

### Behavioral results of the EEG experiment

Figure 2B shows the average amusement rating and negativity rating across the three sentence types in the EEG experiment. For the amusement rating, a one-way repeated-measures ANOVA revealed a significant main effect of sentence type (*F* [2, 70] = 159.58, *p* < 0.001, *η*^*2*^ = 0.82). Pairwise comparison with Bonferroni correction revealed that pun-humor sentences (M = 6.88, SD = 1.24) were significantly more amusing than both non-humor (M = 3.61, SD = 1.08; *p* < 0.001) and nonsensical (M = 4.19, SD = 1.18; *p* < 0.001) sentences, and nonsensical sentences were significantly more amusing than non-humor sentences (*p* < 0.001).

For the negativity rating, a one-way repeated-measures ANOVA revealed a significant main effect of sentence type (*F* [2, 70] = 39.76, *p* < 0.001, *η*^*2*^ = 0.53). Pairwise comparison with Bonferroni correction revealed that pun-humor sentences (M = 7.21, SD = 0.77) were significantly more negative than both non-humor (M = 6.62, SD = 0.84; *p* < 0.001) and nonsensical (M = 6.36, SD = 0.83; *p* < 0.001) sentences, and non-humor sentences were significantly more negative than nonsensical sentences (*p* = 0.022).

### ERP results

Figure 2C displays the grand-averaged ERP waveforms of three sentence types in nine ROIs, and Figure 2D shows the topographic maps of the N400 and LPC components.

### The N400 component (300–500 ms)

The mean amplitudes during 300–500 ms of each ROI were computed and used as the dependent variables. Likelihood-ratio tests indicated that the main effects of Sentence Type (*x*^2^(2) = 19.42, *p* < 0.001) and ROI (*x*^2^(8) = 381.40, *p* < 0.001) were significant. And the interaction between Sentence Type and ROI (*x*^2^(16) = 28.71, *p* = 0.026) provided a better fit for the data than a model without it.

The Bonferroni-corrected pairwise comparisons revealed that compared with nonsensical sentences, negative words in pun- humor sentences elicited significantly smaller N400 components in all ROIs except for the left frontal ROI (*ps* < 0.05). Moreover, N400 components elicited by negative words in pun-humor sentences and non-humor sentences were comparable across all ROIs, except for a marginally significantly smaller N400 component in the medial parietal ROI for pun-humor sentences (*p* = 0.068) (see Table 2 for more details).

**Table 2.**
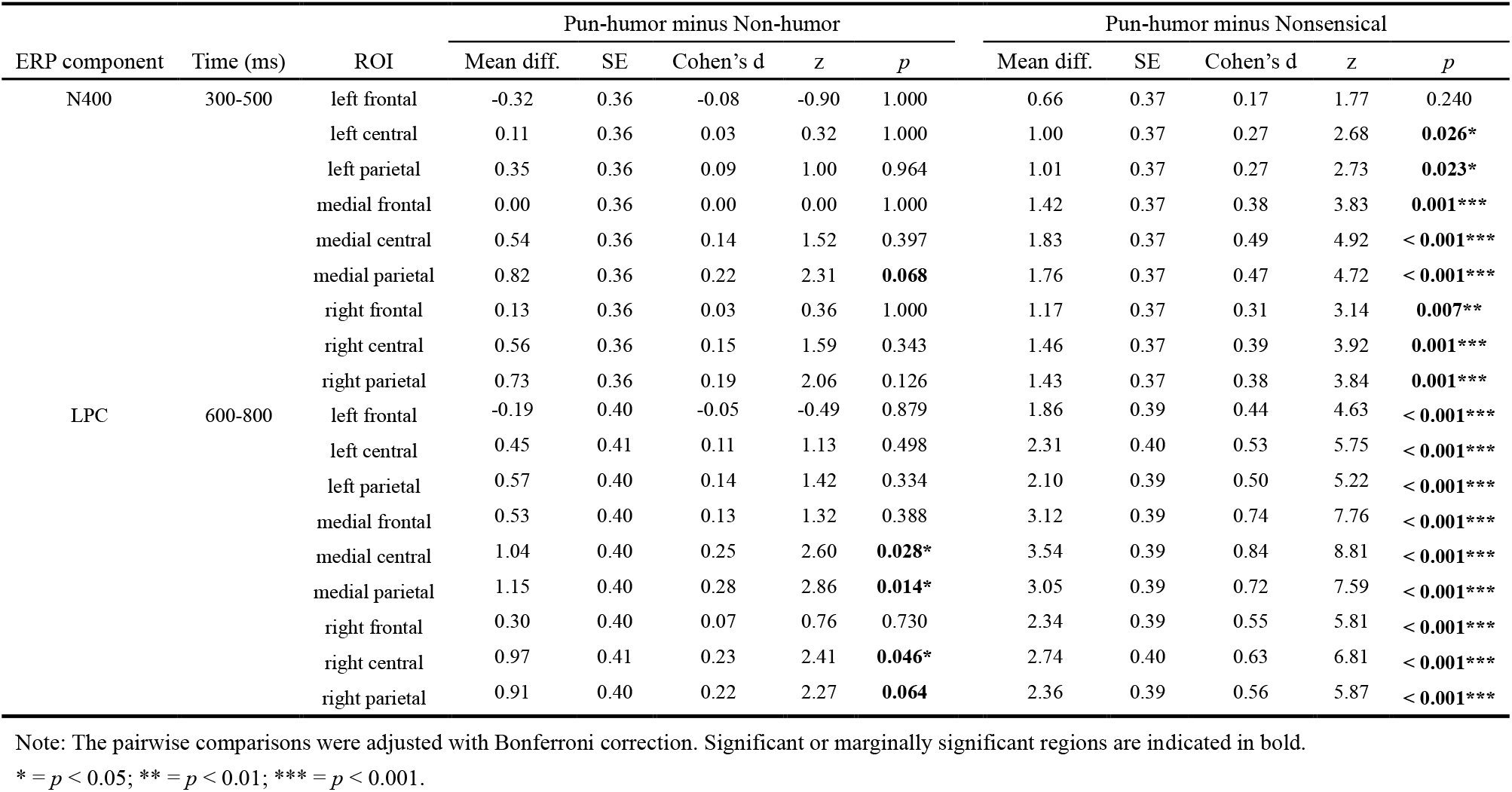
Pairwise comparisons between pun-humor sentences and the other two sentence types for each ROI in two time windows.

### The LPC component (600–800 ms)

The mean amplitudes during 600–800 ms of each ROI were computed and used as the dependent variables. Likelihood-ratio tests indicated that the main effects of Sentence Type (*x*^2^(2) = 45.16, *p* < 0.001) and ROI (*x*^2^(8) = 392.21, *p* < 0.001) were significant. And the interaction between Sentence Type and ROI (*x*^2^(16) = 40.98, *p* < 0.001) provided a better fit for the data than a model without it.

The Bonferroni-corrected pairwise comparisons revealed that compared with nonsensical sentences, negative words in pun- humor sentences elicited significantly greater LPC amplitudes in all ROIs (*ps* < 0.001). Moreover, compared with non-humor sentences, negative words in pun-humor sentences elicited significantly greater LPC amplitudes in medial central (*p* = 0.028), medial parietal (*p* = 0.014), and right central (*p* = 0.046) ROIs, while marginally significantly greater in the right parietal ROI (*p* = 0.064) (see Table 2 for more details).

### RSA results

Regarding the relative timing of representing amusement and negativity in pun-humor sentences, we compared the onset, peak, and duration latencies of the time clusters during which the behavioral patterns of amusement or negativity showed significant partial correlations with the neural patterns. Figure 3A illustrates the RDMs constructed from behavioral data in pun-humor sentences, and Figure 3B shows the RDMs constructed from EEG data at each time point in pun-humor sentences. The RSA results revealed significant partial correlations between the EEG RDMs and behavioral RDMs, indicating the time course of the representation for negativity (Figure 3C) and amusement (Figure 3D) in pun-humor sentences. The representation of negativity began at 230 ms and ended at 670 ms after keyword onset, which specifically comprised three significant clusters: the first spanning from 230 ms to 370 ms (average *p* = 0.008), the second occurring from 400 ms to 540 ms (average *p* = 0.012), and the third ranging from 610 ms to 670 ms (average *p* = 0.027). And the representation of amusement had only one significant cluster, which began at 430 ms and continued until 520 ms (average *p* = 0.025). That is, the representation of negativity lasts 340 ms, which is longer than the 90 ms duration for the representation of amusement. Besides, the representation for negativity peaked around 280 ms and the representation of amusement peaked around 510 ms. For direct comparison, we plotted the intervals of significant time clusters for partial correlations between EEG RDMs and behavioral RDM for negativity and amusement on a single graph (Figure 3E(a)), and averaged the partial correlation values within the significant time clusters to more intuitively reveal the experiential pattern of mixed feelings (Figure 3E(b)). Furthermore, the Bootstrap test showed that the representation of negativity occurred significantly earlier (Figure 3F(a), *p* = 0.015), peaked significantly earlier (Figure 3 F(b), *p* = 0.018), and lasted significantly longer (Figure 3F(c), *p* = 0.007) than amusement in pun-humor sentences.

**Figure 3.**
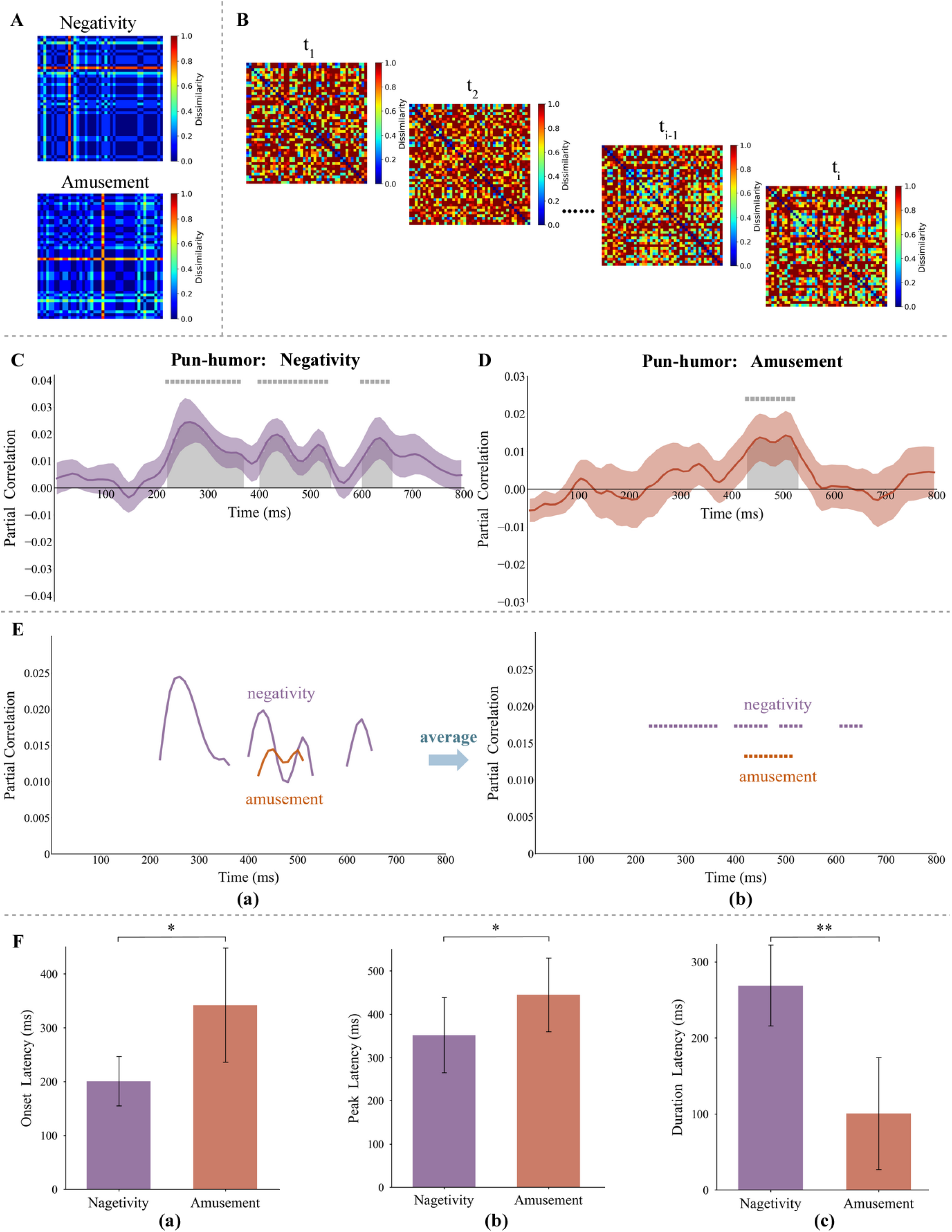
Results of the representational similarity analysis (RSA). A, The two behavioral representational dissimilarity matrixes (RDM) for negativity and amusement. B, The RDMs for EEG data at each time point. C, Time course of partial correlations between EEG RDMs and behavioral RDM for negativity in pun-humor sentences. D, Time course of partial correlations between EEG RDMs and behavioral RDM for amusement in pun-humor sentences. E, (a) Plot the intervals of significant time clusters for partial correlations between EEG RDMs and behavioral RDM for negativity and amusement; (b) average the partial correlation values within the significant time clusters to more intuitively reveal the experiential pattern of mixed emotions. F, Onset (a), peak (b), and duration (c) latencies for decoding negativity and amusement in pun-humor sentences. Considering the large sample size in the bootstrap test, error bars indicate standard deviation. * = *p* < 0.05; ** = *p* < 0.01; *** = *p* < 0.001.

RSA were also conducted on other sentence types to draw the time course of amusement (Figure S2A) and negativity (Figure S2B). As significant partial correlations for negativity representation were observed among the three sentence types, we compared the onset, peak, and duration of the time clusters where behavioral patterns of negativity showed significant partial correlations with neural patterns. This was done to explore the timing of negativity representation across the three sentence types. Specifically, the representation of negativity in non-humor sentences began at 300 ms and ended at 400 ms (average *p* = 0.015), and the representation of negativity in nonsensical sentences comprised two significant clusters: one was 220 ms to 260 ms (average *p* = 0.009), and the other was 490 to 520 ms (average *p* = 0.021). Furthermore, the two-sided bootstrap test with FDR corrected indicated that the onset of representing negativity (Figure S2C(a)) in non-humor sentences was significantly delayed relative to that in pun-humor sentences (*p* = 0.039) and nonsensical sentences (*p* = 0.016), while the onset in pun-humor sentences was comparable to that in nonsensical sentences (*p* = 0.059). Additionally, the duration of representing negativity (Figure S2C(b)) in nonsensical sentences was shorterthan that in non-humor sentences (*p* = 0.026), with pun-humor sentences exhibiting the longest duration (*ps* = 0.023) among all sentence types.

## Discussion

Pun-humor is one of the most important forms of humor, thought to bring positive emotions to people. However, the use of negative keywords in pun-humor sentences can induce mixed feelings (i.e., the co-occurrence of amusement and negativity), where the temporal dynamics of its constituent emotional states during this online processing—the patterns and relative timing of representing amusement and negativity—remain enigmatic. In the present study, behavioral and EEG experiments were conducted and three types of sentences (pun-humor, non-humor, and nonsensical) were used as stimuli to investigate this issue. The behavioral ratings in the two experiments revealed that pun-humor sentences consistently had higher scores in both negativity and amusement ratings as compared to non-humor and nonsensical sentences, suggesting that pun-humor can evoke stronger amusement—which are relatively more positive—yet fails to alleviate and even tends to amplify negativity. And the reading time results showed that participants took longer to process negative words in pun-humor sentences as compared to non-humor sentences, which implies that their processing might require more effort. The ERP results showed that negative keywords in pun-humor sentences elicited comparable N400 components to those in non-humor sentences, but larger LPC components compared to those in non-humor and nonsensical sentences, indicating that keywords in pun-humor sentences can be linked to the context and require increased cognitive effort for more extensive semantic and emotional processing. Notably, the RSA illustrated that the dynamic representation of mixed feelings during the online processing of negative keywords in pun-humor sentences aligned with the highly simultaneous pattern. In this pattern, although the representation of negativity occurred and peaked significantly earlier than amusement, there was a period during which both were simultaneously represented. To the best of our knowledge, this study represents the first attempt to explore the online cognitive and affective processing of negative words in pun-humor sentences, thereby advancing our comprehension of pun-humor’s multifaceted role in social communication. Importantly, this study offers new insights into how the human brain dynamically represents mixed feelings and reveals the neural time courses of different emotional states, which also supports the serial/cascade model of language processing from a novel perspective on emotion processing.

The first goal of the present study is to investigate whether the perceived amusement and negativity counteract each other or are jointly intensified during the real-time processing of pun-humor sentences with negative keywords. At the behavioral level, it was found that pun-humor can elicit stronger positive emotions (i.e., amusement) than non-humor. This result is consistent with the findings in previous studies (e.g., Chan et al., 2012; Feng et al., 2014; Ku et al., 2017), and the stronger positive emotions (i.e., amusement) associated with pun-humor has been commonly regarded as the main reason why humor is effective in regulating negative emotions (e.g., Mobbs et al., 2003; Bartolo et al., 2006; Strick et al., 2009). However, our findings revealed that pun-humor sentences received the highest negativity scores in both the behavioral and EEG experiments, challenging the common assumption that amusement and negativity would counteract each other. One possible explanation for this distinct outcome lies in the methodology employed, specifically the different presentation orders of the stimuli. In most previous studies (e.g., Samson & Gross, 2012; Kugler & Kuhbandner, 2015; Wu et al., 2019), negative stimuli were initially presented, and subsequently, humorous materials emerged as an offline reprocessing form of retroactive adjustment or post-regulation strategy for negative stimuli. In this case, the subsequent humorous information may receive more cognitive resources, leading to potential distraction from the preceding negative stimuli and reducing the perceived negativity (Braniecka et al., 2019; Brawer & Amir, 2021; Chan et al., 2023). However, in the present study, the contexts that can produce punning effects were presented first, followed by the negative words, allowing for the online processing of negative stimuli through humor. Consequently, more cognitive resources are allocated to negative words, which may increase the perceived negativity. The intensified negativity observed in this study is crucial for highlighting the potential adverse effects of employing humor inappropriately to cope with negative stimuli. Currently, there is limited research on the negative impact of humor (Papousek et al., 2023). Although humor is commonly regarded as a complex higher-order emotional process (Van Dillen & Koole, 2007; Farkas et al., 2021; Ruiz-Padial et al., 2023), it is generally believed to have primarily positive implications (e.g., Deckman & Skolnick, 2020; Liao et al., 2023; Erduran Tekin, 2024), as it can make others laugh (Vrticka et al., 2013). Therefore, further investigation is necessary to facilitate a more comprehensive understanding of how humor influences negative stimuli and to raise awareness of its potential negative effects.

The second goal of the present study is to examine how amusement and negativity are dynamically experienced over time during the online processing of pun-humor sentences with negative keywords, which specifically involves the patterns and relative timing of representing amusement and negativity. RSA was utilized to track and analyze the interplay between the inherent negative emotionality of the keywords and the amused nature of pun-humor. It can be observed that the representation of amusement is brief, and within this short duration, the representation of negativity does not disappear but rather persists over a longer time frame. This RSA result corresponds to the behavioral results where pun-humor, despite inducing a strong sense of amusement (Froehlich & Madipakkam,2021; Gloor et al., 2021), cannot diminishes but rather intensifies the negative feelings. Notably, during the processing of negative words in pun-humor, participants experience mixed feelings characterized by a genuine coexistence of both strongly elicited negativity and amusement, rather than the intuitive ‘zero-sum’ process of amusement offsetting negativity. This suggests that the experience of mixed feelings induced by processing pun-humor with negative keywords aligns more closely with the highly simultaneous pattern proposed by Oceja and Carrera (2009). Moreover, the true existence of mixed feelings suggests that emotion categories should not be conceived in a black-or-white thinking manner as solely positive or negative, because this would imply that amusement and negativity cannot occur simultaneously. And the findings of this study reveal that pun-humor embedded with negative keywords is an effective method for exploring the intriguing topic of mixed feelings, offering a valuable source of stimuli that warrants further investigation. Such linguistic material, by simply setting up two language elements (keywords and context) that are homophonic or polysemous yet initially appear inappropriate or unexpected, can effectively elicit the experience of mixed feelings in participants.

Moreover, the relative timing of the onset of specific emotional states linked to different language elements is noteworthy, as it involves not only the processing of mixed feelings but also debates about language processing. In this situation, the onset of negativity is related to the lexical semantic processing of negative keywords, while the onset of amusement is linked to connecting keywords with contexts and uncovering their humorous pragmatic meanings in sentences. Thus, the essence of the question regarding which comes first in the representation of negativity and amusement is to investigate whether accessing the humorous pragmatic meaning is a slow or rapid process. To preliminarily explore this issue, the N400 and LPC components were also compared in this study to verify the cognitive mechanism underlying the online processing of negative keywords in pun-humor. For the N400 component, in the central and parietal regions, negative words in pun-humor sentences elicited comparable and even marginally significantly smaller N400 amplitudes compared to those in non-humor sentences. As for nonsensical sentences, they disrupt the connection by altering a character or word within the context of pun-humor sentences, resulting in processing difficulties in linking keywords with contexts, as evidenced by their largest N400 component across the three sentence types. Similar results were reported in previous studies (Dholakia et al., 2016; Zheng & Wang, 2023a), suggesting that pun-humor sentences may have a potential contextual facilitation effect due to the ability of their keywords to be readily linked with the context in a unique way. Specifically, while the keywords in pun-humor sentences do not directly align semantically with the context as they do in non-humor sentences, they can still be effortlessly connected to the context through homophony or polysemy. And if the keywords have already shown a connection with the context during the N400 time window, it can be speculated that the onset timing of amusement might occur before or within this time window. Our RSA results indeed show that the representation of amusement starts around 400 ms, which coincides roughly with the time window of the N400 component. Therefore, the N400 component represents the formation of connection between negative keywords and their contexts, and it is within this time window that we grasp the humorous pragmatic semantic meaning by cleverly linking these two seemingly unrelated language elements in pun-humor sentences, leading to the emergence of amusement representation.

Notably, negativity representation begins around 200 ms, which is significantly earlier than amusement. When encountering negative keywords in pun-humor sentences, their inherent negativity immediately evokes a negative response in receivers, making it conceivable that the representation of negativity commences as soon as their lexical semantic information is accessed. In prior psycholinguistic studies, it has been discovered that we can access lexical semantics almost “within an instant” (within 200 ms), a process that occurs quickly. Similar results have also been reported in pun-humor studies. For example, a previous eye-tracking study (Zheng et al., 2020) employing the visual world paradigm reported that after the presentation of a pun word, participants displayed a greater fixation on the word associated with its lexical semantic meaning within the first 200 ms. Meanwhile, the onset of negativity significantly precedes amusement, indicating that the access to pragmatic information related to amusement occurs in a later stage than the lexical semantic information associated with negativity, rather than both being processed synergistically at an early stage during online comprehension. This provides novel supporting evidence for the sequential/cascade model in language processing from the perspective of emotional processing, More importantly, the representation of negativity not only onsets and peaks early but also persists for a long period, which even occurrs in the late stages of language processing, within roughly the time window of the LPC component. As for the LPC results, negative keywords were found to elicit significantly larger LPC components in pun-humor sentences than in non-humor and nonsensical sentences, indicating that more cognitive effort is required for more extensive semantic processing of negative keywords in pun-humor sentences (e.g., Coulson & Van Petten, 2002; Rataj et al., 2018; Canal et al., 2019). More specifically, participants need to integrate dual information of negative keyword in pun-humor sentence: one derived from lexical semantic information of the negative keyword itself, and the other from humorous pragmatic information supported by the connection between the keyword and its context. However, in non-humor and nonsensical sentences, only the lexical semantic information needs to be integrated, making the semantic processing less extensive. Considering that both concepts and emotions are activated together during the processing of negative words (e.g., Bower, 1981; Hatzidaki et al., 2015; Rohr & Abdel, 2018), the extent of semantic processing influences the depth of emotional processing (Zhang et al., 2018). From an emotional processing perspective, an enhanced LPC can also indicate that more intricate emotional processing is concurrently activated during increased semantic processing, as evidenced by previous ERP studies which have linked the enhanced LPC to both enhanced late language integration and top-down emotion evaluation (e.g., Zhang et al., 2018; Kuchinke & Mueller, 2019; Pauligk et al., 2019). Thus, the late negativity representation phase observed in the RSA, which was within the LPC time window, can be attributed to the sustained cognitive engagement needed for a deeper level of semantic and emotional processing, indicating a more comprehensive and detailed analysis of negative keywords in pun- humor sentences during the late processing stage.

In summary, the current study examined the real-time processing of negative words in pun-humor and the dynamic representation of mixed feeling blending amusement and negativity. The results showed that negative words in pun-humor sentences evoked stronger amusement but also amplified negativity, underscoring the potential negative impacts when pun- humor is used inappropriately. This may be because pun-humor sentences involve greater cognitive resource investment for the complex processing and analysis of negative keywords, tending to be perceived as sarcastic, which incurs greater harm and cannot be overshadowed by amusement. This complex processing and analysis of negative keywords is supported by ERP results, which show that although negative keywords in pun-humor can easily connect with the context due to homophonic or polysemous relationships, extensive semantic processing of dual information accompanied by elaborate emotional processing is found in the later stages of processing. And using RSA to further explore this elaborate emotional processing revealed that the representation of negativity persists for a longer duration and occurs significantly earlier than that of amusement. The key finding is the discovery of a period where negativity and amusement truly co-occur simultaneously, providing evidence for the genuine existence of mixed feelings, and revealing that the dynamic representation of mixed feelings aligns with the highly simultaneous pattern. Taken together, when reading the sentence *“What is critical for Jim must be hip-hop, because he is hypocritical,”* the lexical negative information of the keyword (*hypocritical*) initially induces perceived negativity, but when uniquely linked to the context (*hip-hop critical*) via shared phonology, we can derive the humorous pragmatic information and experience amusement. Finally, this study wants to further illustrate that pun-humor with negative keywords is a highly practical topic that is increasingly common in contemporary pop culture but is still underexplored, and it also serves as a feasible method for studying mixed feelings and deserves attention in future research.

## Data availability statement

All data that support the findings of this study are available upon reasonable request by contacting with the corresponding author. In consideration of data protection, a formal data sharing agreement is needed when the data are requested.

## Acknowledgement

We would like to thank Dr. Luis Oceja and Dr. Pilar Carrera for offering valuable feedback and suggestions on this work.

## Reference

Bakker, I., Takashima, A., Van Hell, J. G., Janzen, G., & McQueen, J. M. (2015). Tracking lexical consolidation with ERPs: Lexical and semantic-priming effects on N400 and LPC responses to newly-learned words. Neuropsychologia, 79, 33–41. 10.1016/j.neuropsychologia.2015.10.020

Bartolo, A., Benuzzi, F., Nocetti, L., Baraldi, P., & Nichelli, P. (2006). Humor Comprehension and Appreciation: An fMRI Study. Journal of Cognitive Neuroscience, 18(11), 1789–1798. 10.1162/jocn.2006.18.11.1789

Bates, D., Mächler, M., Bolker, B., & Walker, S. (2015). Fitting Linear Mixed-Effects Models Using lme4. Journal of Statistical Software, 67(1). 10.18637/jss.v067.i01

Bekinschtein, T. A., Davis, M. H., Rodd, J. M., & Owen, A. M. (2011). Why Clowns Taste Amused: The Relationship between Humor and Semantic Ambiguity. Journal of Neuroscience, 31(26), 9665–9671. 10.1523/JNEUROSCI.5058-10.2011

Bower, G. H. (1981). Mood and memory. American Psychologist, 36(2), 129–148. 10.1037/0003-066X.36.2.129

Braniecka A., Hanc M., Wolkowicz I., Chrzczonowicz-Stepien A., Mikolajonek A., & Lipiec M. (2019). Is it worth turning a trigger into a joke? Humor as an emotion regulation strategy in remitted depression. Brain Behavior. 9. e01213. 10.1002/brb3.1213

Brawer, J., & Amir, O. (2021). Mapping the ‘amused bone’: Neuroanatomical correlates of humor creativity in professional comedians. Social Cognitive and Affective Neuroscience, 16(9), 915–925. 10.1093/scan/nsab049

Cacioppo, J.T., & Berntson, G.G. (1994). Relationship between attitudes and evaluative space: A critical review, with emphasis on the separability of positive and negative substrates. Psychological Bulletin, 115, 401–423.

Canal, P., Bischetti, L., Di Paola, S., Bertini, C., Ricci, I., & Bambini, V. (2019). ‘Honey, shall I change the baby? – Well done, choose another one’: ERP and time-frequency correlates of humor processing. Brain and Cognition, 132, 41–55. 10.1016/j.bandc.2019.02.001

Chan, Y.-C., Chou, T.-L., Chen, H.-C., & Liang, K.-C. (2012). Segregating the comprehension and elaboration processing of verbal jokes: An fMRI study. NeuroImage, 61(4), 899–906. 10.1016/j.neuroimage.2012.03.052

Chan, Y., Zeitlen, D. C., & Beaty, R. E. (2023). Amygdala-frontoparietal effective connectivity in creativity and humor processing. Human Brain Mapping, 44(6), 2585–2606. 10.1002/hbm.26232

Coulson, S., & Van Petten, C. (2002). Conceptual integration and metaphor: An event-related potential study. Memory & Cognition, 30(6), 958–968. 10.3758/BF03195780

Deckman, K., & Skolnick, A. (2020). Targeting humor to cope with an unpleasant emotion: Disgust. Current Psychology. 42. 10.1007/s12144-020-00798-x.

Delorme, A., & Makeig, S. (2004). EEGLAB: An open source toolbox for analysis of single-trial EEG dynamics including independent component analysis. Journal of Neuroscience Methods, 134(1), 9–21. 10.1016/j.jneumeth.2003.10.009

Dholakia, A., Meade, G., & Coch, D. (2016). The N400 elicited by homonyms in puns: Two primes are not better than one. Psychophysiology, 53(12), 1799–1810. 10.1111/psyp.12762

Erduran Tekin, Ö. (2024). Coping through humor predicts life satisfaction of teachers working in special education institutions: A quantitative and qualitative study. Current Psychology. 10.1007/s12144-023-05555-4

Farkas, A., Trotti, R., Edge, E., Huang, L., Kasowski, A., Thomas, O., Chlan, E., Granros, M., Patel, K., & Sabatinelli, D. (2021). Humor and emotion: Quantitative meta analyses of functional neuroimaging studies. Cortex. 139. 10.1016/j.cortex.2021.02.023.

Feng, Y.-J., Chan, Y.-C., & Chen, H.-C. (2014). Specialization of neural mechanisms underlying the three-stage model in humor processing: An ERP study. Journal of Neurolinguistics, 32, 59–70. 10.1016/j.jneuroling.2014.08.007

Ford, T. E., & Ferguson, M. A. (2004). Social Consequences of Disparagement Humor: A Prejudiced Norm Theory. Personality and Social Psychology Review, 8(1), 79–94. 10.1207/S15327957PSPR0801_4

Friederici, A.D., (2002). Towards a neural basis of auditory sentence processing. Trends in Cognitive Sciences. 6 (2), 78–84. 10.1016/S1364-6613(00)01839-8

Friederici, A.D., (2011). The brain basis of language processing: from structure to function. Psychological Reviews. 91 (4), 1357–1392. 10.1152/physrev.00006.2011

Fritz, H. L., Russek, L. N., & Dillon, M. M. (2017). Humor Use Moderates the Relation of Stressful Life Events With Psychological Distress. Personality and Social Psychology Bulletin, 43(6), 845–859. 10.1177/0146167217699583

Froehlich, E., Madipakkam, A. R., Craffonara, B., Bolte, C., Muth, A. & Park, S. (2021). A short humorous intervention protects against subsequent psychological stress and attenuates cortisol levels without affecting attention. Scientific Reports. 11. 10.1038/s41598-021-86527-1.

Gao, P., Jiang Zh., Yang Y., Zheng Y., Feng G. *, & Li X. * (2024) Temporal neural dynamics of understanding communicative intentions from speech prosody. NeuroImage. 10.1016/j.neuroimage.2024.120830

Gibson, C., & Sagarin, B. J. (2023). Pun-intentionally sadistic: Is punning a manifestation of everyday sadism? Personality and Individual Differences, 203, 111997. 10.1016/j.paid.2022.111997

Gloor, Jamie & Cooper, Cecily & Bowes-Sperry, Lynn & Chawla, Nitya. (2021). Risqué Business? Interpersonal Anxiety and Humor in the #MeToo Era. Journal of Applied Psychology. 107. 10.1037/apl0000937

Goel, V., & Dolan, R. J. (2001). The functional anatomy of humor: Segregating cognitive and affective components. Nature Neuroscience, 4(3), 237–238. 10.1038/85076

Han, T., Xiu, L., & Yu, G. (2020). The impact of media situation on people’s memory effect—An ERP study. Computers in Human Behavior, 104, 106180. 10.1016/j.chb.2019.106180

Hatzidaki, A., Baus, C., & Costa, A. (2015). The way you say it, the way I feel it: Emotional word processing in accented speech. Frontiers in Psychology, 6. 10.3389/fpsyg.2015.00351

Hubbard, R. J., & Federmeier, K. D. (2021). Representational Pattern Similarity of Electrical Brain Activity Reveals Rapid and Specific Prediction during Language Comprehension. Cerebral Cortex, 31(9), 4300–4313. 10.1093/cercor/bhab087

Kato, M., Okumura, T., Tsubo, Y., Honda, J., Sugiyama, M., Touhara, K., & Okamoto, M. (2022). Spatiotemporal dynamics of odor representations in the human brain revealed by EEG decoding. Proceedings of the National Academy of Sciences, 119(21), e2114966119. 10.1073/pnas.2114966119

Kiat, J. E., Hayes, T. R., Henderson, J. M., & Luck, S. J. (2022). Rapid Extraction of the Spatial Distribution of Physical Saliency and Semantic Informativeness from Natural Scenes in the Human Brain. The Journal of Neuroscience, 42(1), 97–108. 10.1523/JNEUROSCI.0602-21.2021

Koleva, K., Mon-Williams, M., & Klepousniotou, E. (2019). Right hemisphere involvement for pun processing – Effects of idiom decomposition. Journal of Neurolinguistics, 51, 165–183. 10.1016/j.jneuroling.2019.02.002

Ku, L.-C., Feng, Y.-J., Chan, Y.-C., Wu, C.-L., & Chen, H.-C. (2017). A re-visit of three-stage humor processing with readers’ surprise comprehension, and amusement ratings: An ERP study. Journal of Neurolinguistics, 42, 49–62. 10.1016/j.jneuroling.2016.11.008

Kuchinke, L., & Mueller, C. J. (2019). Are there similarities between emotional and familiarity-based processing in visual word recognition? Journal of Neurolinguistics, 49, 84–92. 10.1016/j.jneuroling.2018.09.001

Kugler, L., & Kuhbandner, C. (2015). That’s not amused! – But it should be: Effects of humorous emotion regulation on emotional experience and memory. Frontiers in Psychology, 6. 10.3389/fpsyg.2015.01296

Kutas, M., & Federmeier, K. D. (2011). Thirty Years and Counting: Finding Meaning in the N400 Component of the Event-Related Brain Potential (ERP). Annual Review of Psychology, 62(1), 621–647. 10.1146/annurev.psych.093008.131123

Larsen J. T., McGraw A. P., Cacioppo J. T. (2001). Can people feel happy and sad at the same time?. Journal of Personality and Social Psychology, 81, 684-696. doi:10.1037/0022-3514.81.4.684

Lau, E. F., Phillips, C., & Poeppel, D. (2008). A cortical network for semantics: (De)constructing the N400. Nature Reviews Neuroscience, 9(12), 920–933. 10.1038/nrn2532

Lenth, R. (2022). emmeans: Estimated marginal means, aka least-squares means (Version 1.8.3) [Computer software]. https://CRAN.R-project.org/package=emmeans

Li, Y., Zhang, M., Liu, S., & Luo, W. (2022). EEG decoding of multidimensional information from emotional faces. NeuroImage, 258, 119374. 10.1016/j.neuroimage.2022.119374

Liao, Y., Lee, M., Sung, Y., & Chen, H. (2023). The Effects of Humor Intervention on Teenagers’ Sense of Humor, Positive Emotions, and Learning Ability: A Positive Psychological Perspective. Journal of Happiness Studies. 24. 1–19. 10.1007/s10902-023-00654-2.

Lu, Z., & Ku, Y. (2020). NeuroRA: A Python Toolbox of Representational Analysis From Multi-Modal Neural Data. Frontiers in Neuroinformatics, 14, 563669. 10.3389/fninf.2020.563669

Marslen-Wilson, W.D., 1987. Functional parallelism in spoken word-recognition. Cognition 25 (1–2), 71–102.

Marslen-Wilson, W., Tyler, L.K., 1975. Processing structure of sentence perception. Nature 257 (5529), 784–786.

Miyamoto Y, Uchida Y, Ellsworth PC. (2010), Culture and mixed emotions: co-occurrence of positive and negative emotions in Japan and the United States. Emotion. 2010:10(3):404–415.

Mobbs, D., Greicius, M. D., Abdel-Azim, E., Menon, V., & Reiss, A. L. (2003). Humor Modulates the Mesolimbic Reward Centers. Neuron, 40(5), 1041–1048. 10.1016/S0896-6273(03)00751-7

Moeller J, Ivcevic Z, Brackett MA, White AE. (2018), Mixed emotions: network analyses of intra-individual co-occurrences within and across situations. Emotion. 2018:18(8):1106–1121.

Murray, R. J., Kreibig, S. D., Pehrs, C., Vuilleumier, P., Gross, J. J., & Samson, A. C. (2023). Mixed emotions to social situations: An fMRI investigation. NeuroImage, 271, 119973. 10.1016/j.neuroimage.2023.119973

Oceja, L., & Carrera, P. (2009). Beyond a Single Pattern of Mixed Emotional Experience: Sequential, Prevalence, Inverse, and Simultaneous. European Journal of Psychological Assessment, 25(1), 58–67. 10.1027/1015-5759.25.1.58

Papousek, I., Rominger, C., Weiss, E. M., Perchtold, C. M., Fink, A., & Feyaerts, K. (2023). Humor creation during efforts to find humorous cognitive reappraisals of threatening situations. Current Psychology, 42(19), 16176–16190. 10.1007/s12144-019-00296-9

Pauligk, S., Kotz, S. A., & Kanske, P. (2019). Differential Impact of Emotion on Semantic Processing of Abstract and Concrete Words: ERP and fMRI Evidence. Scientific Reports, 9(1), 14439. 10.1038/s41598-019-50755-3

Perchtold-Stefan, C. M., Papousek, I., Rominger, C., Schertler, M., Weiss, E. M., & Fink, A. (2020). Humor comprehension and creative cognition: Shared and distinct neurocognitive mechanisms as indicated by EEG alpha activity. NeuroImage, 213, 116695. 10.1016/j.neuroimage.2020.116695

Pfeifer, V. A., & Pexman, P. M. (2023). Mixed and ambiguous emotions can be studied with verbal irony. Cognitive Neuroscience, 14(2), 65–67. 10.1080/17588928.2023.2181320

Pickering, M.J., Garrod, S., (2004). Toward a mechanistic psychology of dialogue. Behavioral and Brain Sciences, 27 (2), 169–190.

Pickering, M.J., Garrod, S., (2013). An integrated theory of language production and comprehension. Behavioral and Brain Sciences, 36 (4), 329c347.

Popal, H., Wang, Y., & Olson, I. R. (2019). A Guide to Representational Similarity Analysis for Social Neuroscience. Social Cognitive and Affective Neuroscience, 14(11), 1243–1253. 10.1093/scan/nsz099

Prenger, M., Gilchrist, M., Hedger, K., Seergobin, K., Owen, A., & MacDonald, P. (2023). Establishing the Roles of the Dorsal and Ventral Striatum in Humor Comprehension and Appreciation with fMRI. The Journal of Neuroscience. 43. JN-RM. 10.1523/JNEUROSCI.1361-23.2023

Pulvermüller, F., Shtyrov, Y., Hauk, O., (2009). Understanding in an instant: neurophysiological evidence for mechanistic language circuits in the brain. Brain and Language. 110 (2), 81–94. 10.1016/j.bandl.2008.12.001

Rataj, K., Przekoracka-Krawczyk, A., & Van Der Lubbe, R. H. J. (2018). On understanding creative language: The late positive complex and novel metaphor comprehension. Brain Research, 1678, 231–244. 10.1016/j.brainres.2017.10.030

Renoult, L., Wang, X., Calcagno, V., Prévost, M., & Debruille, J. B. (2012). From N400 to N300: Variations in the timing of semantic processing with repetition. NeuroImage, 61(1), 206–215. 10.1016/j.neuroimage.2012.02.069

Ritchie, J. B., Bracci, S., & Op De Beeck, H. (2017). Avoiding illusory effects in representational similarity analysis: What (not) to do with the diagonal. NeuroImage, 148, 197–200. 10.1016/j.neuroimage.2016.12.079

Rohr, C. S., Villringer, A., Solms-Baruth, C., Van Der Meer, E., Margulies, D. S., & Okon-Singer, H. (2016). The neural networks of subjectively evaluated emotional conflicts: Evaluative Neural Responses to Conflict. Human Brain Mapping, 37(6), 2234–2246. 10.1002/hbm.23169

Rohr, L., & Abdel Rahman, R. (2018). Loser! On the combined impact of emotional and person-descriptive word meanings in communicative situations. Psychophysiology, 55(7), e13067. 10.1111/psyp.13067

Ruiz-Padial, E., Moreno-Padilla, M., & Reyes del Paso, G.A. (2023). Did you get the joke? Physiological, subjective and behavioral responses to mirth. Psychophysiology, 60(6), e14292. 10.1111/psyp.14292

Russell, J. A. (1980). A circumplex model of affect. Journal of Personality and Social Psychology, 39, 1161–1178.

Russell, J.A., & Carroll, J.M. (1999). Onthebipolarity ofpositive andnegative affect. Psychological Bulletin, 125, 3–30.

Samson, A. C., & Gross, J. J. (2012). Humour as emotion regulation: The differential consequences of negative versus positive humour.

Schimmack, U. (2001). Pleasure, displeasure, and mixed feelings: Are semantic opposites mutually exclusive? Cognition and Emotion, 15(1), 81–97. 10.1080/0269993004200123

Sheridan, H., Reingold, E. M., & Daneman, M. (2009). Using puns to study contextual influences on lexical ambiguity resolution: Evidence from eye movements. Psychonomic Bulletin & Review, 16(5), 875–881. 10.3758/PBR.16.5.875

Singmann, H., Bolker, B., Westfall, J., Aust, F., & Ben-Shachar, M. (2023). afex: Analysis of factorial experiments (Version 1.2.1) [Computer software]. https://CRAN.R-project.org/package=afex

Stanisławski, K., Cieciuch, J. & Strus, W. (2021). Ellipse rather than a circumplex: A systematic test of various circumplexes of emotions. Personality and Individual Differences. 181, 111052. 10.1016/j.paid.2021.111052

Stokes, M. G., Wolff, M. J., & Spaak, E. (2015). Decoding Rich Spatial Information with High Temporal Resolution. Trends in Cognitive Sciences, 19(11), 636–638. 10.1016/j.tics.2015.08.016

Strick, M., Holland, R. W., Van Baaren, R. B., & Van Knippenberg, A. (2009). Finding comfort in a joke: Consolatory effects of humor through cognitive distraction. Emotion, 9(4), 574–578. 10.1037/a0015951

Strijkers, K., Costa, A., Pulvermüller, F., (2017). The cortical dynamics of speaking: lexical and phonological knowledge simultaneously recruit the frontal and temporal cortex within 200 ms. Neuroimage, 163, 206–219. 10.1016/j.neuroimage.2017.09.041

Tomasello, R., Grisoni, L., Boux, I., Sammler, D., Pulvermüller, F. (2022). Instantaneous neural processing of communicative functions conveyed by speech prosody. Cerebral cortex, 32 (21), 4885–4901. 10.1093/cercor/bhab522

Urbach, T. P., DeLong, K. A., Chan, W.-H., & Kutas, M. (2020). An exploratory data analysis of word form prediction during word-by-word reading. Proceedings of the National Academy of Sciences, 117(34), 20483–20494. 10.1073/pnas.1922028117

Vaccaro, A. G., Kaplan, J. T., & Damasio, A. (2020). Bittersweet: The Neuroscience of Ambivalent Affect. Perspectives on Psychological Science, 15(5), 1187–1199. 10.1177/1745691620927708

Vaccaro, A. G., Wu, H., Iyer, R., Shakthivel, S., Christie, N. C., Damasio, A., & Kaplan, J. (2024). Neural patterns associated with mixed valence feelings differ in consistency and predictability throughout the brain. Cerebral Cortex, 34(4), bhae122. 10.1093/cercor/bhae122

Van Dillen, L. F., & Koole, S. L. (2007). Clearing the mind: A working memory model of distraction from negative mood. Emotion, 7(4), 715–723. 10.1037/1528-3542.7.4.715

Vrticka, P., Black, J. M., & Reiss, A. L. (2013). The neural basis of humour processing. Nature Reviews Neuroscience, 14(12), 860–868. 10.1038/nrn3566

Wang, L., Kuperberg, G., & Jensen, O. (2018). Specific lexico-semantic predictions are associated with unique spatial and temporal patterns of neural activity. eLife, 7, e39061. 10.7554/eLife.39061

Wang, S., Zhang, Y., Shi, W., Zhang, G., Zhang, J., Lin, N., & Zong, C. (2023). A large dataset of semantic ratings and its computational extension. Scientific Data, 10(1), 106. 10.1038/s41597-023-01995-6

Wei, W., Huang, Z., Feng, C., & Qu, Q. (2023). Predicting phonological information in language comprehension: Evidence from ERP representational similarity analysis and Chinese idioms. Cerebral Cortex, 33(15), 9367–9375. 10.1093/cercor/bhad209

Willems, R. M. (2023). MA-EM: A neurocognitive model for understanding mixed and ambiguous emotions and morality. Cognitive Neuroscience, 14(2), 51–60. 10.1080/17588928.2022.2132223

Wu, X., Guo, T., Tan, T., Zhang, W., Qin, S., Fan, J. & Luo, J. (2019). Superior emotional regulating effects of creative cognitive reappraisal. NeuroImage. 200, 540–551. 10.1016/j.neuroimage.2019.06.061

Wu, X., Guo, T., Zhang, C., Hong, T.-Y., Cheng, C.-M., Wei, P., Hsieh, J.-C., & Luo, J. (2021). From “Aha!” to “Haha!” Using Humor to Cope with Negative Stimuli. Cerebral Cortex, 31(4), 2238–2250. 10.1093/cercor/bhaa357

Xu, Q., Hu, J., Qin, Y., Li, G., Zhang, X., & Li, P. (2023). Intention affects fairness processing: Evidence from behavior and representational similarity analysis of event-related potential signals. Human Brain Mapping, 44(6), 2451–2464. 10.1002/hbm.26223

Zhang, M., Ge, Y., Kang, C., Guo, T., & Peng, D. (2018). ERP evidence for the contribution of meaning complexity underlying emotional word processing. Journal of Neurolinguistics, 45, 110–118. 10.1016/j.jneuroling.2016.07.002

Zhang, J., Wu, C., Yuan, Z., and Meng, Y. (2018). Different early and late processing of emotion-label words and emotion-laden words in second language: an ERP study. Second Language Research. 36, 399–412. 10.1177/0267658318804850

Zhendong Dong & Qiang Dong. (2003). HowNet—A hybrid language and knowledge resource. International Conference on Natural Language Processing and Knowledge Engineering, 2003. Proceedings. 2003, 820–824. 10.1109/NLPKE.2003.1276017

Zheng, W., & Wang, X. (2022). Contextual Support for Less Salient Homophones and Pun Humor Appreciation: Evidence From Eye Movements in Reading Chinese Homophone Puns. Frontiers in Psychology, 13, 875479. 10.3389/fpsyg.2022.875479

Zheng, W., & Wang, X. (2023a). Frame-shifting instead of incongruity is necessary for pun comprehension: Evidence from an ERP study on Chinese homophone puns. Language, Cognition and Neuroscience, 1–14. 10.1080/23273798.2023.2192509

Zheng, W., & Wang, X. (2023b). Humor experience facilitates ongoing cognitive tasks: Evidence from pun comprehension. Frontiers in Psychology, 14, 1127275. 10.3389/fpsyg.2023.1127275

Żochowska, A., Nowicka, M. M., Wójcik, M. J., & Nowicka, A. (2021). Self-face and emotional faces—Are they alike? Social Cognitive and Affective Neuroscience, 16(6), 593–607. 10.1093/scan/nsab020

