## supplementary_information for "Processing negative words in pun-humor: dynamic representation of mixed feelings blending amusement and negativity"

Table 1. Pairwise comparisons between pun-humor sentences and the other two sentence types for each ROI in baseline intervals.

| | Time (ms) | ROI | Pun-humor minus Non-humor | | | | | Pun-humor minus Nonsensical | | | | |
|---|---|---|---|---|---|---|---|---|---|---|---|---|
| | | | Mean diff. | SE | Cohen's d | z | p | Mean diff. | SE | Cohen's d | z | p |
| Baseline | [-200, 0] | left frontal | 0.20 | 0.48 | 0.03 | 0.42 | 1.000 | 0.21 | 0.42 | 0.04 | 0.49 | 1.000 |
| | | left central | -0.20 | 0.48 | 0.03 | -0.43 | 1.000 | -0.11 | 0.42 | 0.02 | -0.26 | 1.000 |
| | | left parietal | -0.37 | 0.48 | 0.06 | -0.78 | 1.000 | -0.34 | 0.42 | 0.07 | -0.79 | 1.000 |
| | | medial frontal | -0.44 | 0.48 | 0.07 | -0.92 | 1.000 | -0.14 | 0.42 | 0.03 | -0.35 | 1.000 |
| | | medial central | -0.65 | 0.48 | 0.11 | -1.36 | 0.520 | -0.51 | 0.42 | 0.10 | -1.20 | 0.688 |
| | | medial parietal | -0.09 | 0.48 | 0.02 | -0.19 | 1.000 | -0.53 | 0.42 | 0.10 | -1.25 | 0.628 |
| | | right frontal | -0.30 | 0.48 | 0.05 | -0.61 | 1.000 | -0.57 | 0.42 | 0.11 | -1.32 | 0.552 |
| | | right central | -0.50 | 0.48 | 0.09 | -1.02 | 0.920 | -0.28 | 0.42 | 0.05 | -0.57 | 1.000 |
| | | right parietal | -0.71 | 0.48 | 0.12 | -1.47 | 0.421 | -0.41 | 0.42 | 0.08 | -0.96 | 1.000 |

Note: The pairwise comparisons were adjusted with Bonferroni correction. There are no significant regions.

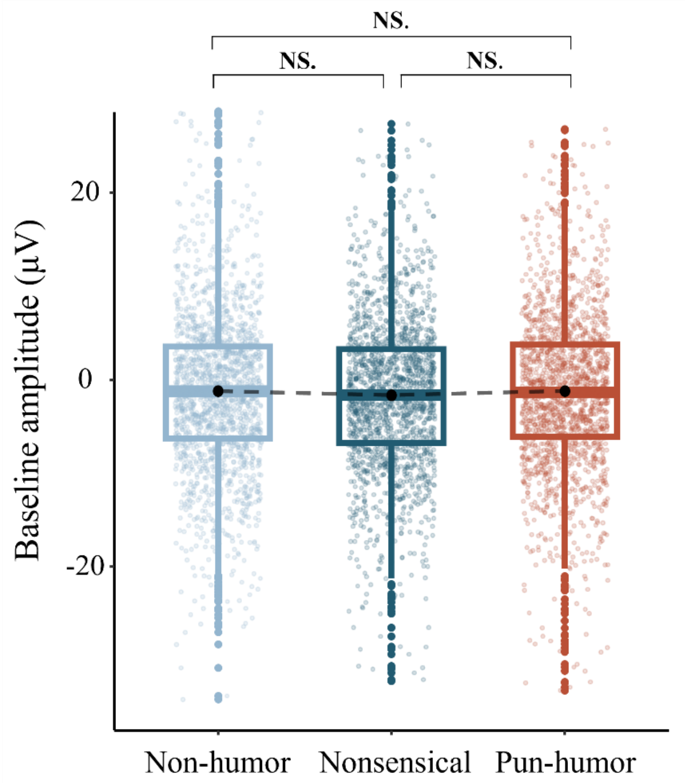


**Figure S1.** The amplitudes during baseline intervals (200-0 ms before keyword onset) among three sentence types showed no significant differences.

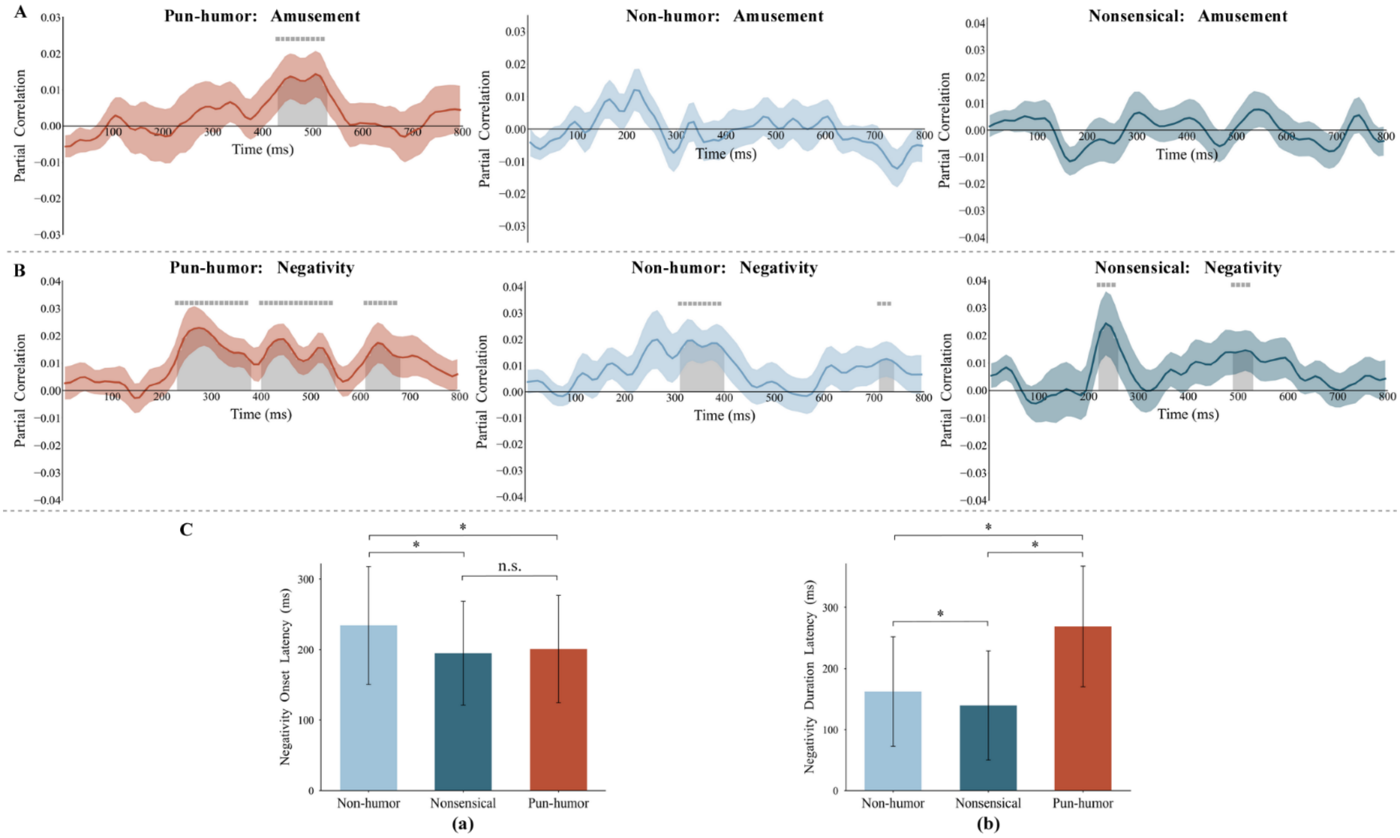


**Figure S2.** A, Time course of partial correlations between EEG RDMs and behavioral RDM for amusement in pun-humor, non-humor, and nonsensical sentences. B, Time course of partial correlations between EEG RDMs and behavioral RDM for negativity in pun-humor, non-humor, and nonsensical sentences. C, Onset (a), and duration (b) latencies for decoding negativity in three types of sentences.
